# Colonization resistance against *Clostridioides difficile* is a graded, microbiota-intrinsic property of healthy human gut communities

**DOI:** 10.64898/2026.08.26.747331

**Authors:** Gurjit Sidhu, Daniel Marquina, Tara Share, Joan Whitlock, Jennifer Gollwitzer, Ajisha Alwin, James Martin, Gary P. Wang

## Abstract

Fecal microbiota transplantation cures approximately 90% of recurrent *Clostridioides difficile* infection, yet it remains unknown whether all healthy donor microbiota confer equivalent protection. We colonized germ-free C57BL/6 mice with stool microbiota from 30 healthy human donors and challenged them with *C. difficile* in the absence of antibiotic pretreatment. Donor microbiota conferred a spectrum of colonization resistance phenotypes: Resistant (no detectable colonization or toxin), Carrier (asymptomatic colonization with detectable toxin), Symptomatic (non-lethal diarrheal illness), and Susceptible (lethal infection). Of these, 8 conferred Resistant phenotypes, 12 Carrier, 6 mixed Resistant–Carrier outcomes, and 4 Symptomatic or Susceptible phenotypes. While 16S rRNA gene sequencing of donor stool did not distinguish phenotypes across any diversity or compositional metric tested, humanized mouse microbiomes exhibited clear phenotype-dependent differences after engraftment. Richness (observed amplicon sequence variants) and diversity (Shannon and Faith’s phylogenetic diversity) declined progressively from Resistant to Susceptible phenotypes, although substantial overlap was observed between groups. Differential abundance analysis identified taxa depleted across non-resistant phenotypes, including Lachnospiraceae taxa such as *Hungatella* and *Sellimonas*, and *Bacteroides intestinalis*. Shotgun metagenomics confirmed these associations and revealed coordinated depletion of biosynthetic and carbohydrate metabolism pathways in non-resistant phenotypes, consistent with broad loss of community metabolic capacity rather than loss of a single dominant function. These findings demonstrate colonization resistance is a graded, microbiota-associated ecological property, evident after host engraftment rather than being a binary trait encoded in donor stool. This has implications for donor screening in fecal microbiota transplantation and the rational design of microbiome-based therapeutics.

**Lay Summary:** Clostridioides difficile is a leading cause of serious diarrhea in hospitals. Fecal microbiota transplantation, which transfers stool from a healthy donor to a patient, can cure many recurrent infections. This strongly suggests that healthy gut microbes can protect against this pathogen. However, it is unclear whether all healthy donors provide the same level of protection.

In this study, we colonized germ-free mice with gut microbial communities from thirty healthy adult donors and then challenged them with Clostridioides difficile. The donor communities produced a range of outcomes: some completely blocked the pathogen, some allowed asymptomatic carriage, some caused mild diarrhea, and a few led to severe disease. Importantly, the make-up of donor stool alone did not predict these outcomes. Differences only became apparent after the microbes had established themselves in the mouse gut. Protection was associated with higher overall microbial diversity, certain bacterial groups, and a broad range of metabolic functions rather than any single species.

These findings show that protection is not a fixed property of donor stool, but emerges from how the microbial community establishes and functions in the host. These results have implications for improving donor selection and developing more effective microbiome-based therapies.

## Introduction

*Clostridioides difficile* is the leading cause of healthcare-associated infectious diarrhea [1], responsible for substantial morbidity, mortality, and healthcare costs worldwide [2,3]. Antibiotic exposure is the primary risk factor [1], with advanced age and hospitalization further increasing susceptibility [4].

Fecal microbiota transplantation (FMT) achieves high cure rates (>90% with repeat administration) for recurrent *C. difficile* infection [5], providing strong evidence that the gut microbiota plays a central role in colonization resistance [6]. Proposed mechanisms include bile acid metabolism, short-chain fatty acid production, nutrient competition, and direct microbial antagonism [7]. Such insights have driven the development of defined live biotherapeutic products, such as spore-enriched formulations [8] and defined commensal consortia [9], to replace empiric fecal transplantation with rationally selected microbial consortia.

FMT and live biotherapeutic development commonly assume that microbiota from healthy donors protect against *C. difficile.* However, donor screening typically excludes pathogens and clinical risk factors rather than directly assessing colonization resistance potential [10,11]. Most experimental models also rely on antibiotic pretreatment to induce host susceptibility [12,13], thereby obscuring the transplanted microbial community’s intrinsic colonization resistance capacity. Studies using germ-free mice colonized with human microbiota have either used pooled rather than individual donor communities [14], or challenged individual donor-colonized mice without antibiotics. They found *C. difficile* colonized across donors without clear donor-dependent resistance [15], which leaves unresolved whether individual healthy donor communities differ in their ability to confer resistance.

Here, we address this question by colonizing germ-free mice with microbiota from 30 individual healthy donors and challenging them with *C. difficile* in the absence of antibiotic perturbation. We show that microbiota from individual healthy donors confer a spectrum of colonization resistance phenotypes and that these phenotypes are not predictable from donor stool composition alone. Using complementary 16S rRNA gene sequencing and shotgun metagenomics, we identify taxonomic and functional features associated with variation across this spectrum of outcomes.

## Methods

### Donor recruitment

The study was approved by the University of Florida Institutional Review Board (IRB #202000542). Thirty healthy adults were recruited between March and December 2021 to provide fresh stool samples (**Table 1**). Donors were screened to exclude those with active or recent gastrointestinal symptoms or antibiotic use within the preceding six months.

**Table 1:** Characteristics of healthy stool donors (N=30)

|  |  |
| --- | --- |
| Age, years: median (IQR) | 27 (22.5) |
| Sex |  |
| Male | 15 (50%) |
| Female | 15 (50%) |
| Race |  |
| White | 19 (63%) |
| Asian | 5 (17%) |
| Black or African American | 6 (20%) |
| Body Mass Index (BMI): median (IQR) |  |
| Male BMI | 25.8 (4.6) |
| Female BMI | 25.8 (11.7) |

### Preparation of donor stool gavage

Stool samples were collected in Raku-Ryu flushable cups, transferred to 50 mL sterile collection tubes (Corning, cat#430897), and transported to the laboratory within 2 hours of defecation. All subsequent steps were performed under anaerobic conditions within 2 hours of receipt. A 10% w/v suspension was prepared in pre-reduced phosphate-buffered saline (PBS), vortexed for 10 minutes, and centrifuged at 500g for 10 minutes at room temperature. The supernatant was aliquoted into microcentrifuge tubes (Eppendorf Protein Lo-Bind, #0030108442) and stored at −80°C.

### *Clostridioides difficile* culture

*Clostridioides difficile* VPI 10463 spore stocks were serially diluted in 1% bovine serum albumin (BSA) to prevent clumping and plated anaerobically onto taurocholate cefoxitin cycloserine fructose agar (TCCFA). After 48 hours, colonies were counted and gavage inocula were prepared at 1,000 spores per 100 µL in PBS, aliquoted, and stored at 4°C. Spore counts were confirmed by plating each inoculum onto TCCFA after challenge.

### Mouse Experiments

All animal experiments were approved by the University of Florida Institutional Animal Care and Use Committee (IACUC #202009773). Individually housed, sex-matched germ-free (GF) C57BL/6 mice (6–10 weeks old) were kept in HEPA-filtered NexGen cages with autoclaved food and water in a dedicated ABSL2 suite.

Mice were orally gavaged with 100 µL of donor stool suspension to establish colonization. After a 21-day colonization period, humanized mice and germ-free controls were challenged with 1,000 spores of *C. difficile* VPI 10463 by oral gavage. Pre-challenge fecal pellets were collected for 16S rRNA gene sequencing to confirm colonization. Post-challenge, mice were weighed and assessed daily for up to 14 days using a standardized health scoring system (Supplemental Table S1). Mice were euthanized upon reaching humane endpoints (sustained weight loss >15% or moribund condition); surviving mice were euthanized on day 14 post challenge.

### Murine cecum and colon collection

Mice were immediately necropsied after euthanasia. Cecal and colonic contents were collected into microcentrifuge tubes and stored on ice. The cecum and colon were flushed with PBS, bisected longitudinally, and then swiss-rolled. Tissues were fixed in 10% formalin for 24 hours, transferred to 70% ethanol, and stored at 4°C until processing by the University of Florida Molecular Pathology Core.

### *C. difficile* load quantification

Cecal contents were transferred to an anaerobic chamber, suspended in pre-reduced PBS, and plated onto TCCFA. Colonies were counted after 48-hour anaerobic incubation at 37°C.

### 16S rRNA V1-V3 amplicon sequencing

DNA was extracted from mouse fecal pellets using the DNeasy PowerSoil Pro Kit (Qiagen, #47016) and from human stool using the PSP Spin Stool DNA Basic Kit (Invitek Diagnostics, Germany, #1038120300) with Stool DNA Stabilizer tubes (Invitek Diagnostics, Germany, #1038111100) per manufacturer’s instructions. The V1–V3 hypervariable region of the 16S rRNA gene was amplified using barcoded primers 27F (5′-AGAGTTTGATCCTGGCTCAG-3′) and 534R (5′-ATTACCGCGGCTGCTGG-3′) in 20 µL reactions containing 4 µL template, 100 nM of each primer, and 10 µL SuperFi PCR master mix (Invitrogen). Amplicons were verified by 1% SYBR Safe agarose gel electrophoresis, gel-purified using the NucleoSpin Gel and PCR Clean-Up Kit (Qiagen, Valencia, CA, USA, #740609) and quantified using the Qubit HS DNA quantification kit (Invitrogen, Carlsbad, CA, USA). Equimolar amplicons were pooled, library concentration was determined by qPCR (Library Quant Kit, Cat#KK4824, Roche Diagnostics), and the multiplexed library was sequenced on an Illumina MiSeq using the MiSeq Reagent Kit v3 (600-cycle, Illumina, San Diego, CA, USA).

### Bioinformatics

MiSeq reads were demultiplexed using Cutadapt4 [16] and imported into QIIME2 (v2024.10) [17]. Reads were denoised and amplicon sequence variants were called using the DADA2 pipeline [18] via q2-dada2 plugin, and taxonomy was assigned to ASVs using the q2-feature-classifier [19] classify-sklearn naive Bayes taxonomy classifier against the SILVA v138 [20] reference database. Phylogenetic trees were generated using the QIIME2 phylogeny plugin. Alpha and beta diversity metrics were calculated using the QIIME2 diversity plugin. Differential abundance analysis was performed using ANCOM-BC [21] via qiime composition ancombc plugin. Graphs and statistical analyses were generated in GraphPad Prism v10.1 or R (v4.5, [22].

### Shotgun Metagenomics

Shotgun libraries were prepared from pre-challenge fecal DNA using the Illumina DNA Prep kit (20060060) and sequenced on a NovaSeq X (10B kit) at the ICBR NextGen DNA Sequencing Core at University of Florida. Reads were processed using KneadData (v 0.12.0): low-quality bases and adapter sequences were trimmed using Trimmomatic (v 0.39) [23], and host (mouse) reads were removed by mapping against the mouse_C57BL_6NJ reference genome using Bowtie2 [24]. Cleaned paired-end reads were concatenated per sample. Taxonomic profiles were generated using MetaPhlAn4 (v 4.1.1) [25], and gene and pathway abundances were estimated using HUMAnN3 (v 3.8) [26]. Differential abundance analysis was performed using MaAsLin3 [27] or LefSe [28].

## Results

### Healthy donor microbiota confer a spectrum of colonization resistance phenotypes

We colonized germ-free C57BL/6 mice with stool microbiota from 30 individual donors and challenged them with 1,000 spores of *C. difficile* VPI 10463 after a 21-day colonization period (**Figure 1A**) to determine whether microbiota from healthy donors differ in intrinsic colonization resistance. This inoculum was uniformly lethal in germ-free controls. Among 167 humanized mice, 137 remained asymptomatic following challenge, whereas 30 developed clinical signs including diarrhea, weight loss, or moribund decline. Of these, 20 survived to the 14-day endpoint, while 10 became moribund within four days.

**Figure 1.**
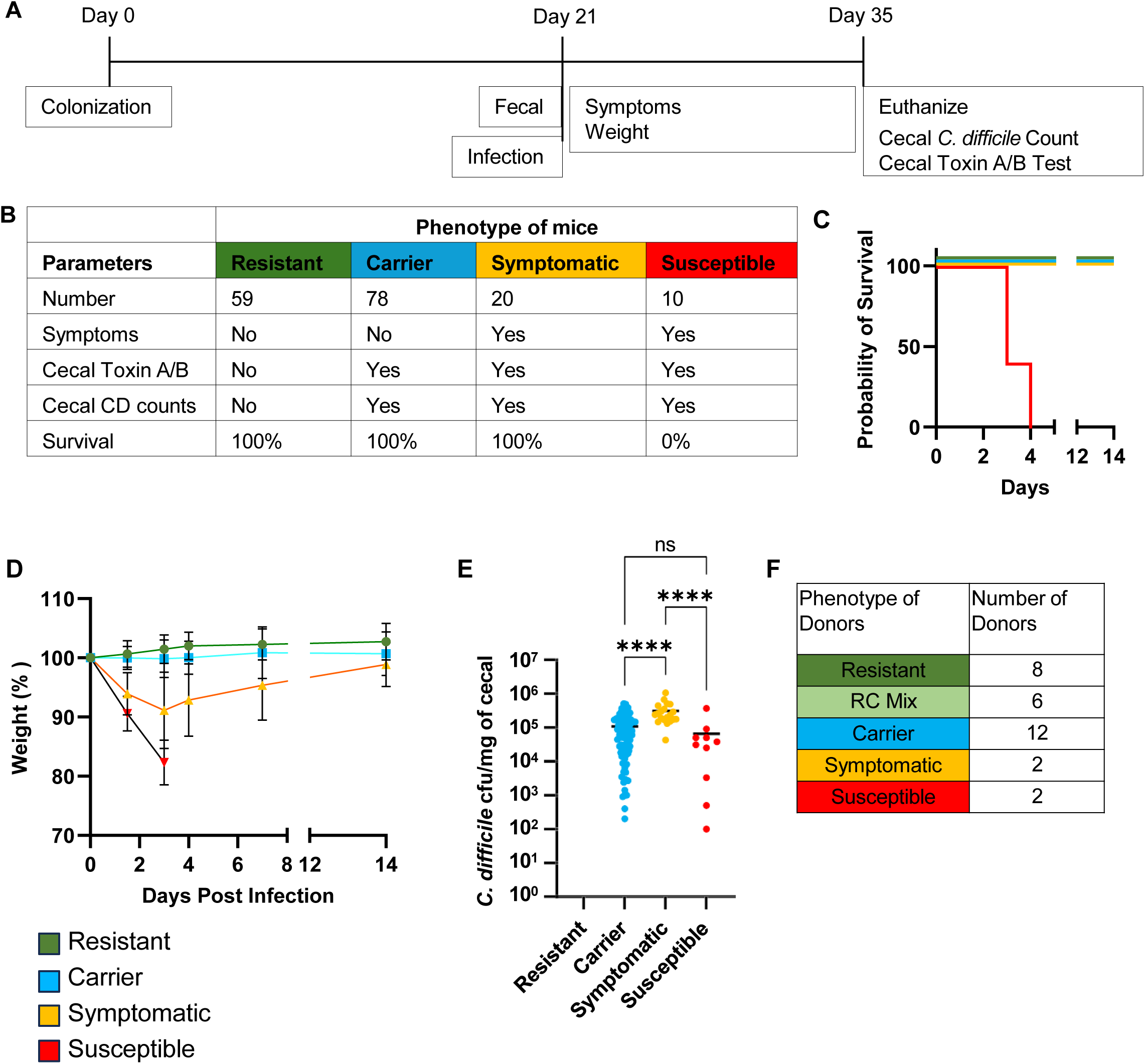
Healthy donor microbiota confer a spectrum of colonization resistance phenotypes in gnotobiotic mice. **(A)** Experimental design. Germ-free C57BL/6 mice were colonized with individual donor stool microbiota on day 0. Pre-challenge fecal samples were collected on day 21, followed by oral challenge with 1,000 spores of *C. difficile* VPI 10463. Mice were monitored daily for symptoms and body weight for 14 days post-challenge, with cecal contents collected at endpoint for *C. difficile* quantification and toxin detection. **(B)** Mouse Phenotype classification. Mice were classified into four phenotypes based on clinical symptoms, cecal toxin detection, cecal *C. difficile* burden, and survival. **(C)** Survival curves by phenotype. Susceptible mice (n = 10) became moribund and were euthanized within 4 days post-challenge. Resistant (n = 59), Carrier (n = 78), and Symptomatic (n = 20) mice survived to endpoint. **(D)** Body weight expressed as a percentage of pre-challenge baseline. Data are mean ± SEM. Susceptible mice exhibited rapid weight loss prior to euthanasia. Symptomatic mice showed initial weight loss followed by recovery to baseline by endpoint. **(E)** Cecal *C. difficile* burden at endpoint (CFU/mg cecal contents). Resistant mice had no detectable colonies. ****p < 0.0001; ns, not significant (Ordinary One Way ANOVA). **(F)** Donor-level colonization resistance phenotypes (N = 30). Donor phenotype was assigned based on the predominant outcome (≥80% concordance) among recipient mice (**Supplemental Table S2**). Donors with approximately equal proportions of Resistant and Carrier mice were classified as RC Mix; all recipient mice from these donors survived to endpoint.

Mice were classified into four phenotypes based on clinical outcome, cecal *C. difficile* burden, and toxin detection. Resistant mice had no detectable *C. difficile* or toxin. Carrier mice harbored detectable *C. difficile* and toxin but remained asymptomatic, indicating that toxin detection alone was insufficient to produce overt clinical disease. Symptomatic mice had loose stools throughout the observation period but survived to endpoint, whereas Susceptible mice developed severe disease and required euthanasia (**Figure 1B**). Weight trajectories were consistent with phenotype classification. Resistant and Carrier mice maintained baseline weight, whereas Symptomatic mice exhibited initial weight loss to 91.1% ± 6.4% at 36 hours post-infection, followed by recovery to 98.9% ± 3.7% by endpoint (**Figure 1D**). Cecal *C. difficile* burden in Symptomatic mice (mean 3.1 × 10⁵ CFU/g, n = 20) was approximately threefold higher than in Carrier mice (mean 1.0 × 10⁵ CFU/g, n = 78) (**Figure 1E**). Histopathological scores in the cecum and colon increased progressively with phenotype severity (**Supplemental Figure S1**).

Assignment of donor phenotype based on ≥80% concordance among recipient mice showed that 8 donors (26%) were Resistant, 6 (20%) were Resistant–Carrier mix, 12 (40%) were Carrier, 2 (7%) were Symptomatic, and 2 (7%) were Susceptible (**Figure 1F**; **Supplemental Table S1**). These results demonstrate that microbiota from healthy donors confer a spectrum of colonization resistance phenotypes, from pathogen exclusion (Resistant) through asymptomatic colonization (Carrier) and non-lethal disease (Symptomatic) to lethal susceptibility (Susceptible), rather than conferring uniform protection.

### Donor stool microbiome composition does not predict colonization resistance phenotype

16S rRNA gene sequencing of donor stools showed no significant differences in alpha diversity across phenotype groups, including observed amplicon sequence variants, Pielou’s evenness, Shannon diversity, and Faith’s phylogenetic diversity (**Supplemental Figure S2**). Beta diversity analysis using Jaccard, Bray–Curtis, unweighted UniFrac, and weighted UniFrac distances showed no phenotype-based clustering (**Supplemental Figure S3**), and PERMANOVA confirmed no significant separation between phenotype groups (p = 0.516–0.54; **Supplemental Table S2**). These findings indicate that conventional compositional profiling of donor stool is not predictive of colonization resistance phenotype.

### Humanized mouse microbiomes exhibit phenotype-dependent differences prior to *C. difficile* challenge

Analysis of pre-challenge fecal microbiomes (day 21) demonstrated a progressive decline in richness from Resistant to Carrier to Symptomatic to Susceptible groups. Observed amplicon sequence variants (ASVs) declined significantly across successive phenotype transitions (Resistant > Carrier > Symptomatic; **Figure 2**). Similar trends were observed for Pielou’s evenness, Shannon diversity, and Faith’s phylogenetic diversity. Differences between Symptomatic and Susceptible groups were not statistically significant, likely due to the small number of Susceptible donors (n = 2). Beta diversity analysis showed separation of microbiomes by phenotype across multiple distance metrics (**Figure 3**). PERMANOVA confirmed significant differences between all pairwise phenotype comparisons (p ≤ 0.001; **Supplemental Tables S3A–B**). These findings indicate that phenotype-associated microbiome structure emerges during host engraftment and is not apparent in donor stool composition alone.

**Figure 2.**
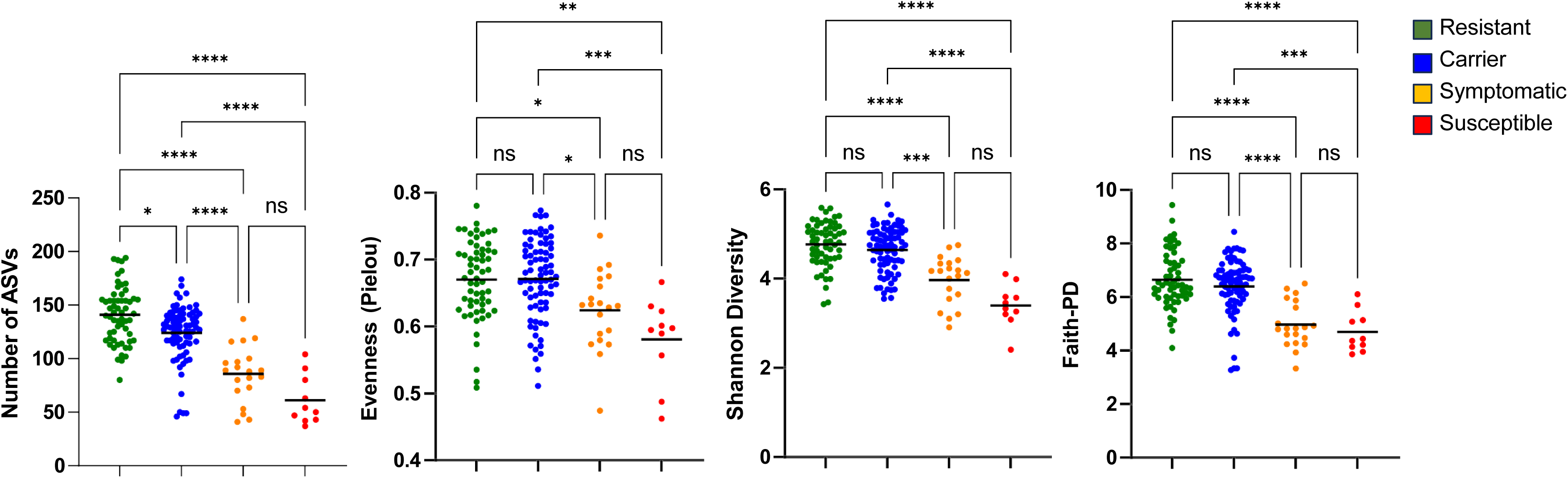
Alpha diversity of pre-challenge fecal microbiomes declines progressively with decreasing colonization resistance. Alpha diversity metrics—observed amplicon sequence variants (ASVs), Pielou’s evenness, Shannon diversity, and Faith’s phylogenetic diversity—are shown for pre-challenge fecal microbiomes collected on day 21 from humanized mice, grouped by colonization resistance phenotype. Horizontal bars indicate means. Richness (observed ASVs) and diversity (Shannon diversity and Faith’s phylogenetic diversity) progressively declined from Resistant to Susceptible phenotypes. Differences between Symptomatic and Susceptible groups were not statistically significant for any metric. Each dot represents one mouse. *p < 0.05, **p < 0.01, ***p < 0.001, ****p < 0.0001; ns, not significant (Kruskal–Wallis test with Dunn’s multiple comparisons).

**Figure 3.**
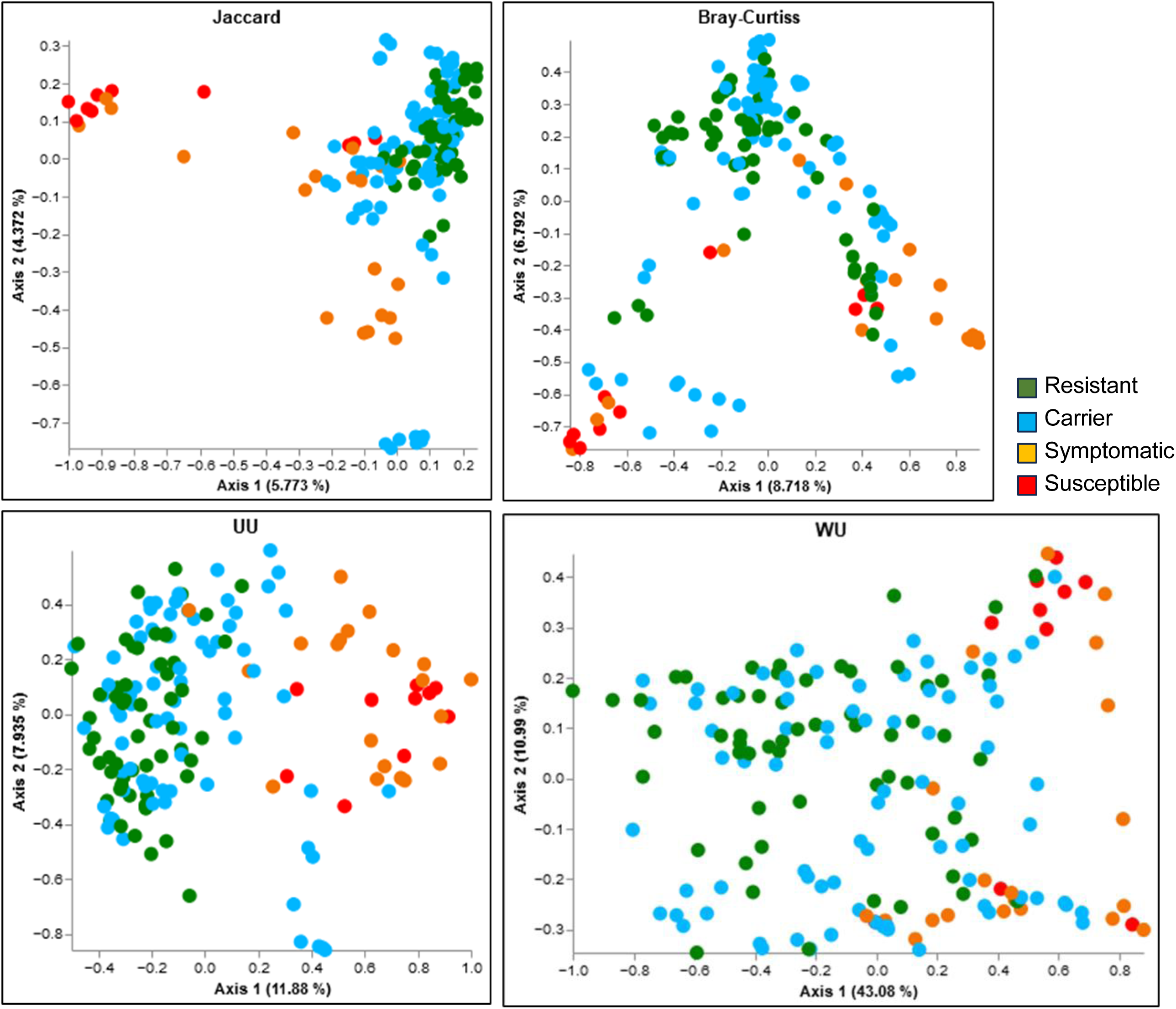
Beta diversity of pre-challenge fecal microbiomes differs by colonization resistance phenotype. Principal coordinates analysis (PCoA) of Jaccard, Bray–Curtis, unweighted UniFrac (UU), and weighted UniFrac (WU) distances among pre-challenge fecal microbiomes from humanized mice, colored by colonization resistance phenotype. Microbiomes demonstrated phenotype-dependent separation across all four distance metrics. Each point represents one mouse. The percentage of variance explained by each axis is indicated on the axis labels. PERMANOVA confirmed significant differences between all pairwise phenotype comparisons (p ≤ 0.001; **Supplemental Tables S4**).

### Endpoint cecal microbiomes maintain phenotype-specific community structure

Richness (observed amplicon sequence variants [ASVs]) decreased across phenotypes, although differences between Resistant and Carrier groups were not statistically significant (**Supplemental Figure S4**). Shannon diversity and Faith’s phylogenetic diversity were significantly lower in Symptomatic microbiomes compared to Resistant and Carrier groups. Beta diversity analysis demonstrated separation of microbiomes by phenotype across Jaccard, Bray–Curtis, and UniFrac distances (**Supplemental Figure S5**). PERMANOVA confirmed significant differences between Resistant, Carrier, and Symptomatic groups (p ≤ 0.001; **Supplemental Table S4A-B**). These findings indicate that phenotype-associated community structure is established prior to infection and persists through the endpoint.

### Richness alone does not determine colonization resistance phenotype

Some Resistant microbiomes exhibited lower richness than Carrier microbiomes, and some Carrier microbiomes exhibited richness comparable to Symptomatic or Susceptible groups (**Supplemental Figure S6**). Similar variability was observed across fecal (**Supplemental Figure S6A**) and cecal (**Supplemental Figure S6B**) microbiomes. These findings indicate that while reduced richness is associated with loss of colonization resistance, richness correlates with, but does not determine, resistance phenotype.

### Colonization of mice by human microbiota is partial and variable

On average, 50% of donor amplicon sequence variants (ASVs) were detected in pre-challenge fecal microbiomes and 55% in endpoint cecal microbiomes (donor stool mean 246 ASVs vs. 124 in fecal and 136 in cecal samples; **Supplemental Figure S7A**). Engraftment efficiency varied considerably across donors (**Supplemental Figure S7B**). This incomplete and variable engraftment suggests that functional resistance phenotypes may arise from selective retention or loss of specific community members during host colonization.

### Specific taxa distinguish transitions between colonization resistance states

Differential abundance analysis identified taxa depleted in Carrier microbiomes across fecal and/or cecal samples, predominantly members of Bacillota, particularly Lachnospiraceae taxa such as *Hungatella* and *Sellimonas*, along with *Bacteroides intestinalis* (**Figure 4A**). Of these, 10 belonged to Bacillota, 2 to Bacteroidota, and 1 to Pseudomonadota. Six taxa were consistently depleted in both fecal and cecal microbiomes: *Bacteroides intestinalis*, Lachnospiraceae sp., *Hungatella* sp., *Sellimonas* sp., *Parabacteroides* sp., and *Catenibacillus* sp. Several of these taxa were further depleted in Symptomatic and Susceptible microbiomes, consistent with progressive loss of specific community members as colonization resistance declined (**Figure 4B**).

**Figure 4.**
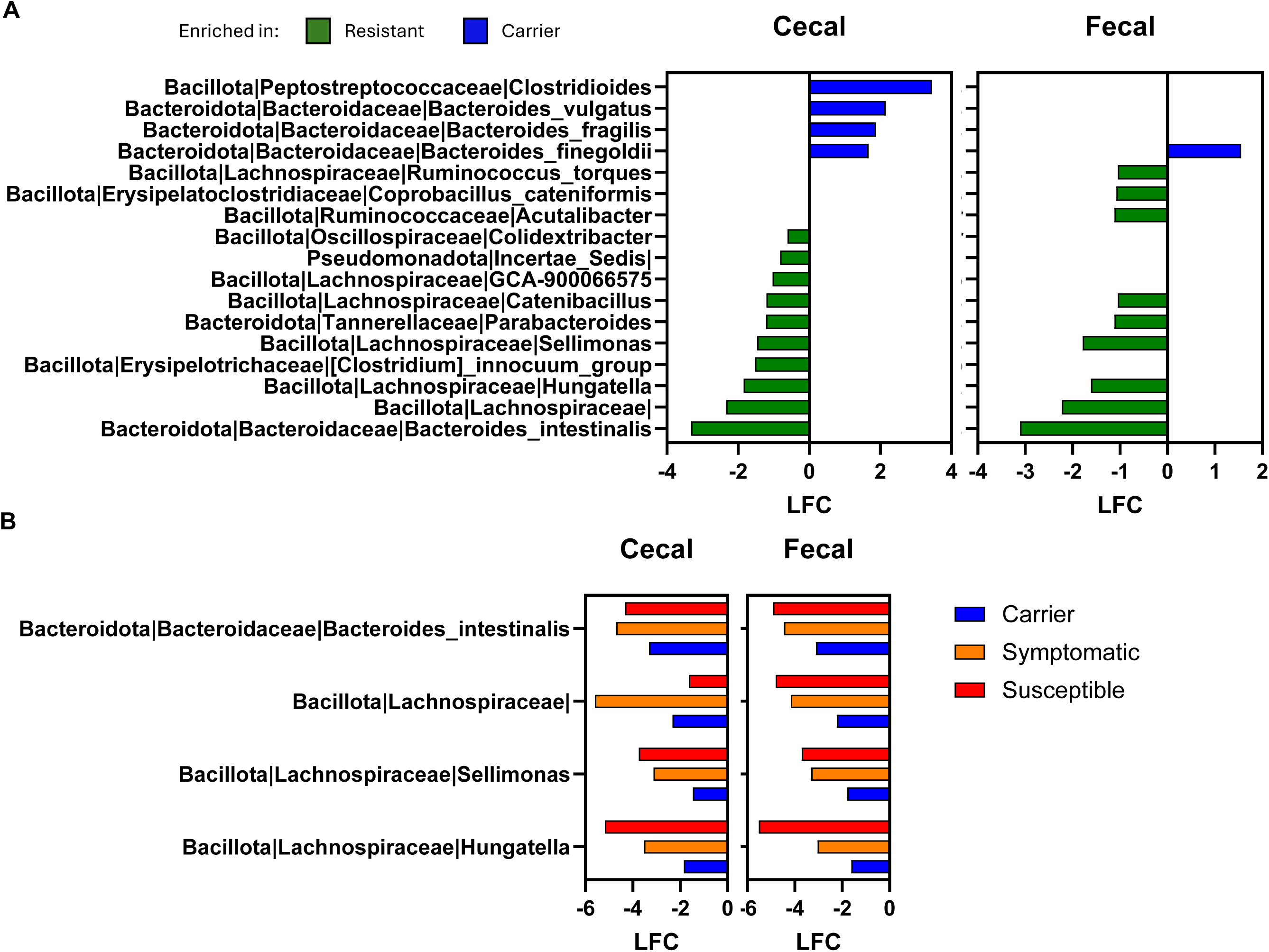
Differential abundance analysis identifies taxa depleted in non-resistant microbiomes. **(A)** Log-fold change (LFC) of taxa in endpoint cecal (left) and pre-challenge fecal (right) Carrier microbiomes relative to Resistant microbiomes. Green bars indicate taxa depleted in Carrier microbiomes (enriched in Resistant), whereas blue bars indicate taxa enriched in Carrier microbiomes (depleted in Resistant). **(B)** Log-fold change of taxa depleted in Carrier, Symptomatic, and Susceptible cecal and fecal microbiomes relative to Resistant microbiomes. Differential abundance analysis was performed using ANCOM-BC (QIIME2 v2024.10) for taxa reaching statistical significance (q<0.05).

Comparison of Carrier and Symptomatic microbiomes identified 18 taxa depleted in endpoint cecal microbiomes and 14 in pre-challenge fecal Symptomatic microbiomes (**Figure 5**). Of the 18 depleted cecal taxa, 13 belonged to Bacillota, 4 to Bacteroidota, and 1 to Pseudomonadota. Of these, fourteen fecal-depleted taxa were also observed to be depleted in cecal microbiomes. These findings support a stepwise ecological model in which progressive loss of specific community members tracks with transitions from resistance to colonization and from colonization to disease.

**Figure 5.**
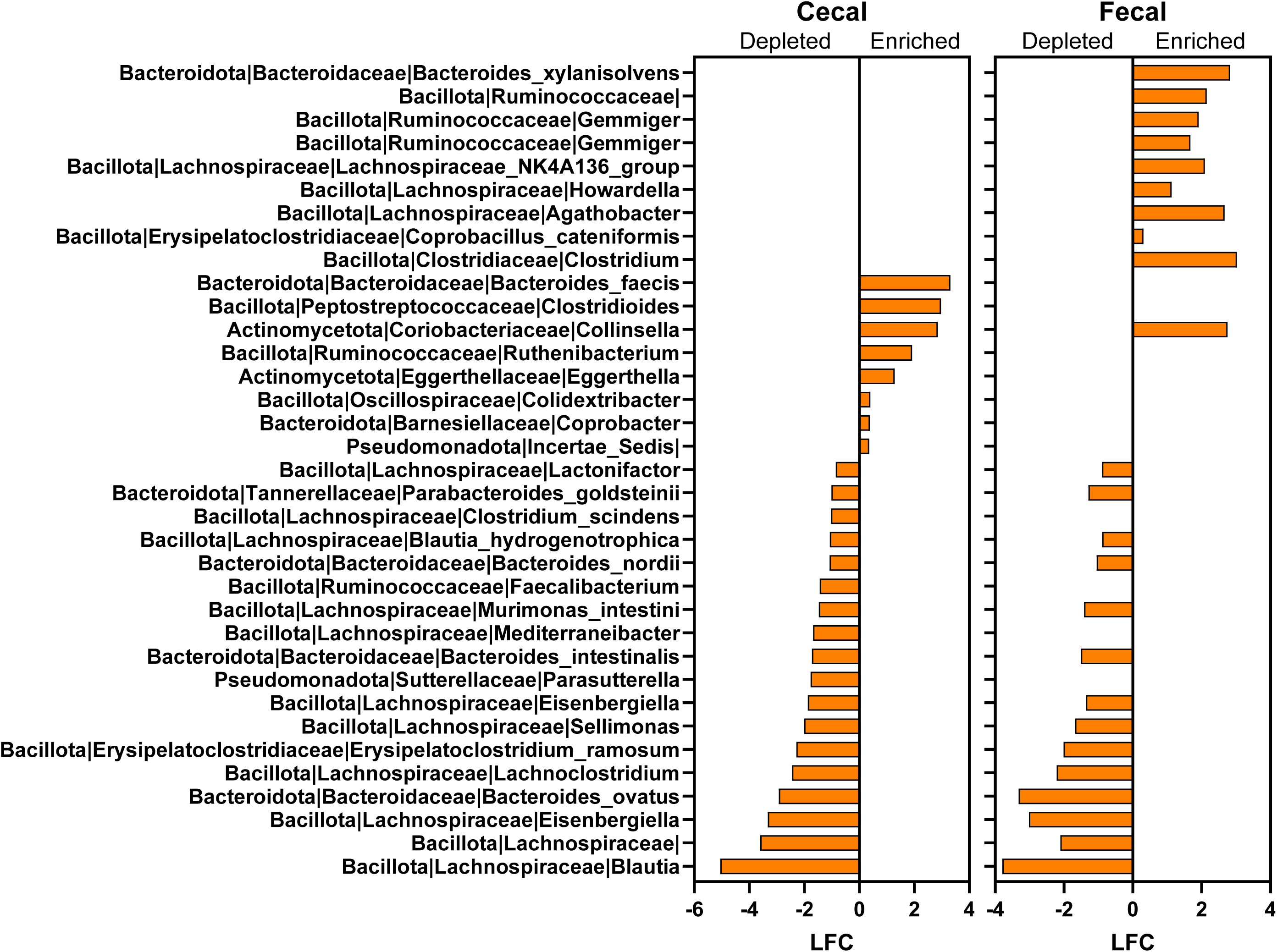
Differential abundance analysis identifies taxa depleted in Symptomatic microbiomes relative to Carrier microbiomes. Log-fold change (LFC) of taxa in endpoint cecal (left) and pre-challenge fecal (right) Symptomatic microbiomes relative to Carrier microbiomes. Bars extending to the left indicate taxa depleted in Symptomatic microbiomes, while those extending to the right indicate enriched taxa. Eighteen taxa were depleted in endpoint cecal and 14 in pre-challenge fecal Symptomatic microbiomes, with 14 common to both. Of the 18 depleted cecal taxa, 13 belonged to Bacillota, 4 to Bacteroidota, and 1 to Pseudomonadota. Seventeen taxa were enriched in Symptomatic microbiomes (8 cecal, 10 fecal), with 1 common to both. Differential abundance analysis was performed using ANCOM-BC (QIIME2 v2024.10). Only taxa reaching statistical significance (q<0.05) are shown.

### Shotgun metagenomics confirms taxonomic and functional differences

Shotgun metagenomic sequencing of pre-challenge fecal microbiomes confirmed taxonomic differences observed by 16S rRNA gene sequencing. Differential abundance analysis identified 8 genera, 12 species, and 15 species-level genome bins enriched in Resistant metagenomes, predominantly within Bacillota and Bacteroidota (**Figure 6**). Notably, *Bacteroides intestinalis*, *Hungatella hathewayi*, and Lachnospiraceae taxa were consistently enriched in Resistant microbiomes across both datasets, providing cross-platform validation.

**Figure 6.**
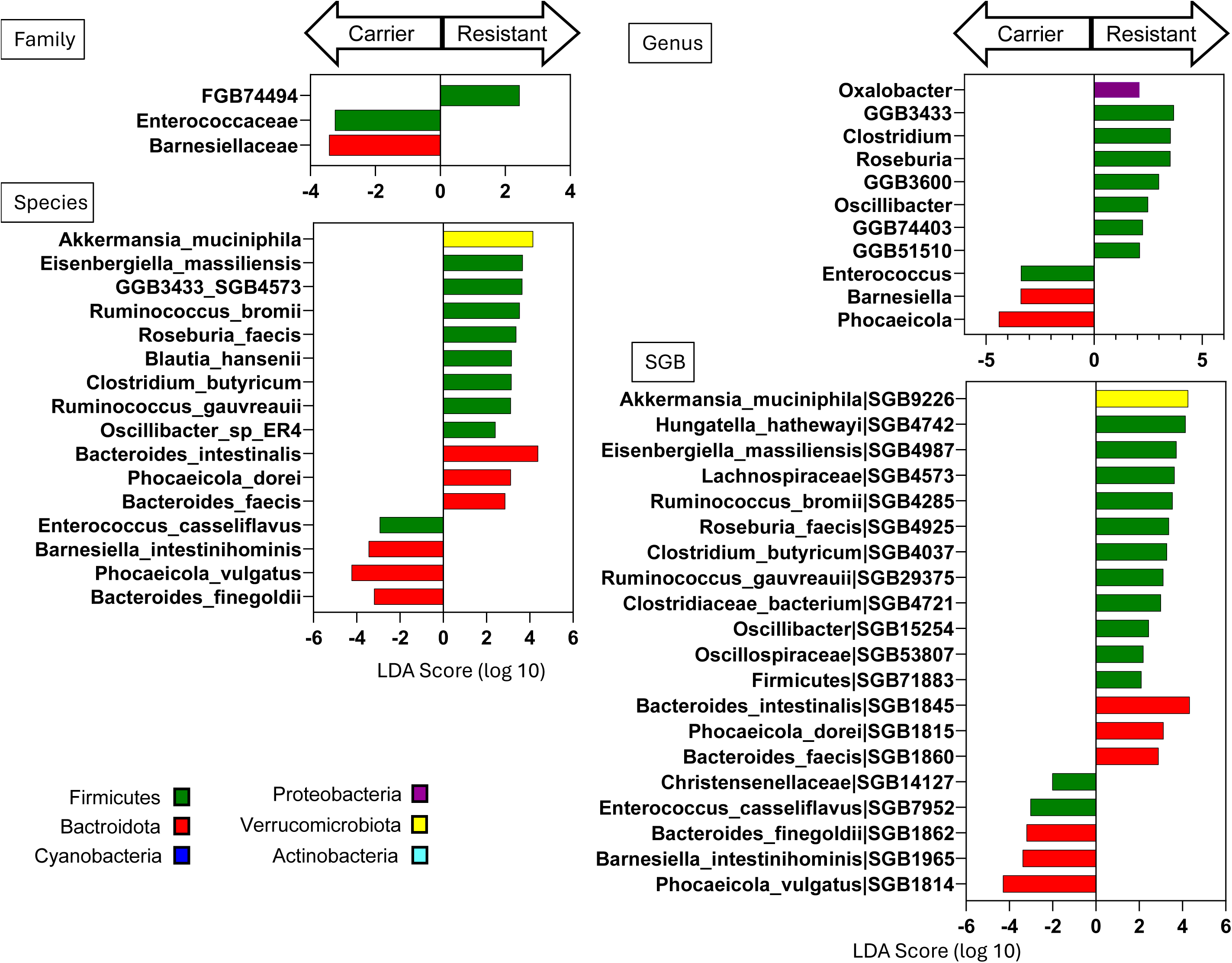
LEfSe analysis identifies taxa enriched in Resistant and Carrier metagenomes. Linear discriminant analysis effect size (LEfSe) analysis was performed on pre-challenge fecal metagenomes from Resistant (n = 8) and Carrier (n = 11) mice at the family, genus, species, and species-level genome bin (SGB) levels. LDA scores (log₁₀) are shown for taxa significantly enriched in Resistant (bars extending to the right) or Carrier (bars extending to the left) metagenomes. Taxa are colored by phylum. Only taxa with an LDA score > 2 and p < 0.05 are shown.

Functional pathway analysis revealed marked differences across colonization resistance phenotypes. A total of 56 pathways were depleted in Susceptible relative to Resistant metagenomes, whereas none were enriched. In Carrier microbiomes, only two pathways were significantly depleted relative to Resistant microbiomes: the pentose phosphate pathway and its non-oxidative branch I (**Figure 7A**). The majority of pathways depleted in Susceptible microbiomes were involved in various biosynthetic processes—including those for amino acids, nucleotide, cofactor, and cell structure—as well as carbohydrate metabolism and energy production (**Figure 7B**; **Supplemental Figure S7**).

**Figure 7.**
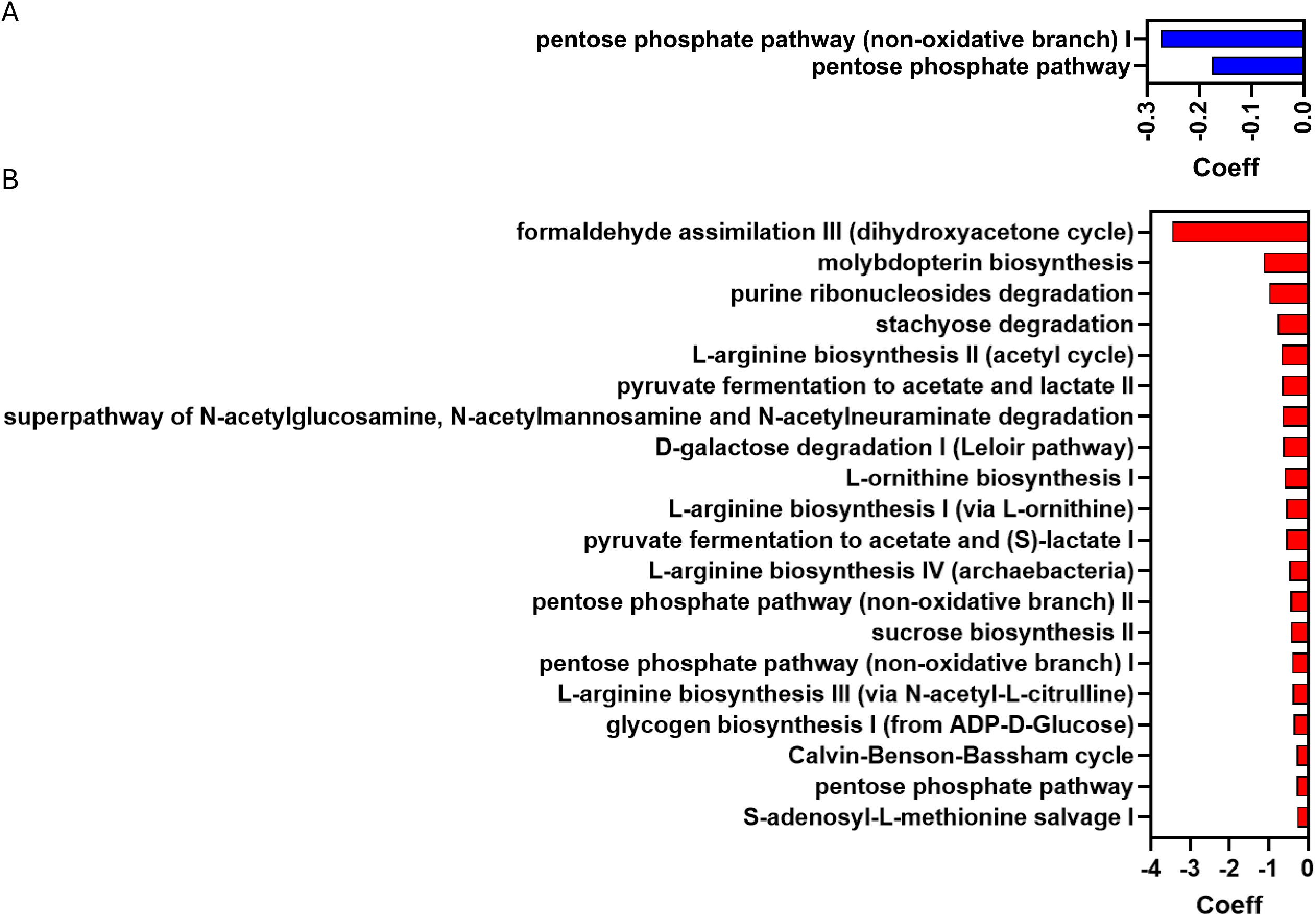
Metabolic pathways depleted in Carrier and Susceptible metagenomes relative to Resistant. **(A)** Pathways significantly depleted in Carrier metagenomes relative to Resistant metagenomes (q < 0.05). The x-axis shows the β-coefficient from MaAsLin3. Two pathways were depleted: the pentose phosphate pathway and its non-oxidative branch I. **(B)** Top 20 pathways depleted in Susceptible metagenomes relative to Resistant metagenomes (q < 0.05). The x-axis shows the β-coefficient from MaAsLin3. The top 20 of 56 depleted pathways are shown, spanning amino acid biosynthesis, carbohydrate degradation, cofactor biosynthesis, and central carbon metabolism.

At the gene family level, 32 MetaCyc reactions were depleted in Carrier and 264 in Susceptible metagenomes relative to Resistant, with 24 of the 264 Susceptible-depleted features completely absent from Susceptible metagenomes (**Figure 8**). These changes were distributed across multiple pathways, consistent with broad loss of community metabolic capacity and reduced functional redundancy, rather than failure of a single dominant protective pathway.

**Figure 8.**
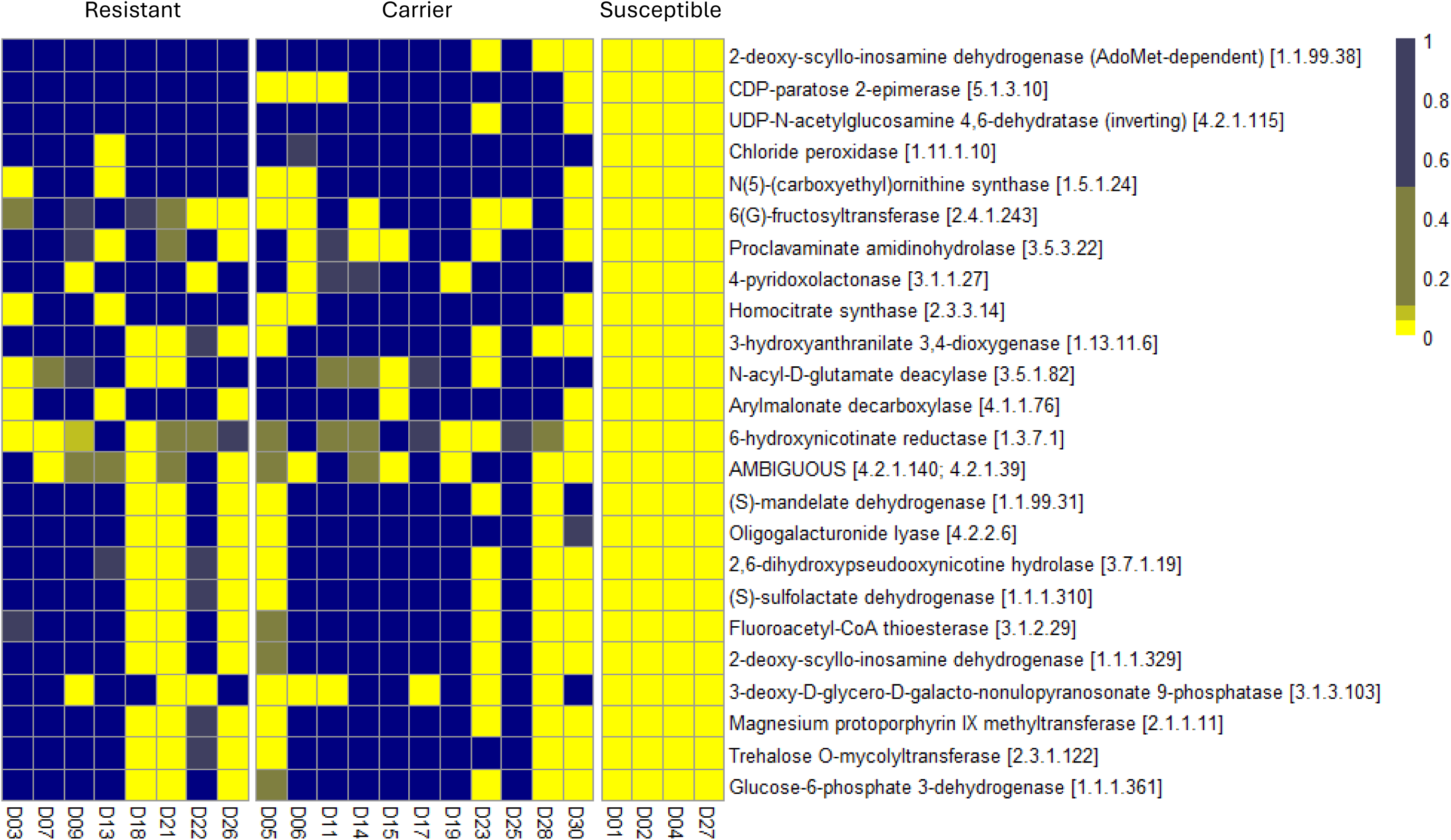
Gene families absent in Susceptible metagenomes but present in Resistant metagenomes. Heatmap showing the relative abundance of 24 gene families (MetaCyc reactions) that are entirely absent (prevalence = 0) in Susceptible metagenomes but present in at least 50% of Resistant metagenomes. Each column represents one donor; donors are grouped by colonization resistance phenotype (Resistant, Carrier, Susceptible) as indicated on the x-axis. Each row represents one gene family, labeled by enzyme name and EC number. The color scale indicates relative abundance (yellow = absent, blue = high); data are not scaled, and the upper range is capped at 1 to enhance visualization of low-abundance differences.

## Discussion

Colonization resistance against *Clostridioides difficile* is a graded, microbiota-intrinsic ecological trait, not a binary property of healthy microbiomes, emerging after host engraftment. By colonizing germ-free mice with microbiota from 30 individual healthy donors in the absence of antibiotic perturbation, we observed a spectrum of outcomes ranging from complete pathogen exclusion to lethal susceptibility. These results define a four-state model of colonization resistance — Resistant, Carrier, Symptomatic, and Susceptible — spanning pathogen exclusion, asymptomatic colonization, non-lethal disease, and lethal infection. In addition, a subset of donors exhibited mixed Resistant– Carrier outcomes across recipient mice, indicating that colonization resistance is graded across donors and that some communities may reside near a functional threshold at which minor variation in engraftment composition shifts the outcome between adjacent phenotypic states.

A central finding of this study is that colonization resistance phenotype was not predictable based on donor stool microbiome composition. Across multiple diversity metrics and compositional analyses, donor microbiota that conferred resistance were indistinguishable from those that permitted colonization or disease. This result has direct implications for current approaches to fecal microbiota transplantation (FMT), which rely on clinical screening and pathogen exclusion rather than functional assessment of microbiome activity [10,11]. Together, these findings challenge the assumption that clinically healthy donor microbiota are functionally equivalent with respect to colonization resistance — an assumption that has also been challenged in other FMT indications, where donor-dependent efficacy has been observed [29].

In contrast, phenotype-associated differences emerged after host engraftment. Humanized mouse microbiomes exhibited progressive declines in richness and diversity across the resistance spectrum, with clear separation by phenotype in beta diversity analyses. These findings indicate that colonization resistance is not encoded in donor composition alone but emerges through host-mediated ecological filtering during engraftment. In this framework, the host environment selects for or against specific microbial configurations, resulting in functionally distinct community states that are not evident from donor stool profiling.

Although reduced diversity was associated with loss of colonization resistance, richness alone did not determine phenotype. Substantial overlap in diversity metrics across groups, including instances of relatively low-diversity Resistant microbiomes and high-diversity Carrier or Symptomatic microbiomes, indicates that global diversity measures alone are insufficient to explain functional outcomes. These observations support a model in which colonization resistance reflects specific community configurations and interactions rather than overall diversity per se.

Consistent with this interpretation, differential abundance analysis identified specific taxa that were progressively depleted across non-resistant phenotypes, primarily within the Bacillota phylum, especially the Lachnospiraceae family. Several taxa, including *Bacteroides intestinalis*, *Hungatella*, and *Sellimonas*, were consistently depleted in Carrier microbiomes and further reduced in Symptomatic and Susceptible states, consistent with a stepwise loss of key community members as resistance declined. While these associations do not establish causality, they are consistent with prior work implicating members of Bacillota and Bacteroidota in colonization resistance through bile acid metabolism, nutrient competition, and production of inhibitory metabolites [7,30,31].

Shotgun metagenomic analysis further demonstrated that functional differences across phenotypes were distributed across multiple pathways rather than driven by a single dominant mechanism. Susceptible microbiomes showed coordinated depletion of pathways involved in biosynthesis, carbohydrate metabolism, and energy production, along with loss of numerous gene families. These findings support a model of colonization resistance as a distributed ecological function arising from metabolic capacity and redundancy within the microbial community, rather than dependence on any single pathway or dominant taxon.

An important feature of this study is the identification of a Carrier phenotype characterized by detectable *C. difficile* colonization and toxin production in the absence of overt disease. This observation suggests that colonization and toxin presence can be decoupled from clinical outcome, a concept supported by prior work demonstrating host modulation of toxin-mediated disease [32], consistent with models in which host-or community-level mechanisms buffer the effects of toxin production. This phenotype may have parallels in human populations, including asymptomatic carriers [33] and settings where colonization may not correlate with disease burden [34][35].

Several limitations should be considered. First, this study was conducted in germ-free mice, and while this model enables controlled assessment of microbiota-intrinsic properties, it does not fully recapitulate the complexity of human host physiology or immune responses. Second, engraftment of human microbiota into mice was incomplete and variable, with approximately 60% of donor ASVs detected in recipient microbiomes, potentially altering community structure and function. Third, the number of donors conferring Symptomatic and Susceptible phenotypes was relatively small, which may limit statistical power for inter-group comparisons. Finally, while taxonomic and functional associations were identified, causal relationships between specific microbes or pathways and resistance phenotypes were not directly tested.

In summary, our findings demonstrate that colonization resistance to *C. difficile* is a graded, emergent ecological property that arises from host–microbiota interactions and reflects distributed functional capacity within the microbial community. These findings have important implications for the design of microbiome-based therapeutics, suggesting that effective interventions may require reconstruction of community-level functional capacity rather than supplementation with individual taxa. More broadly, this work underscores the importance of functionally assessing microbiota, beyond compositional profiling, to predict and understand host–microbe interactions.

## Acknowledgement

This work was supported by NIAID R21 AI150250, VA CSR&D Merit Review I01CX001391, and the Gatorade Trust through funds distributed by the University of Florida, Department of Medicine.

## Data availability statement

The datasets generated during and/or analyzed during the current study are available in the NIH NLM Sequence Read Archive (SRA) repository, under BioProject ID: PRJNA1458299 (https://dataview.ncbi.nlm.nih.gov/object/PRJNA1458299).

## Author contributions

Conceptualization: Gary P. Wang

Funding Acquisition: Gary P Wang

Investigation: DM, TS, JW, JM, GS, JG, AA

Methodology: GPW, GS, JM, DM, AA

Formal Analysis: GS

Project Administration: GS

Resources: GS, JW

Supervision: GPW, GS

## Ethical approval statement

All animal experiments were approved by the University of Florida Institutional Animal Care and Use Committee (IACUC #202009773)

## Patient consent statement

The study was approved by the University of Florida Institutional Review Board (IRB #202000542)

