## Supplemental Material for "Colonization resistance against *Clostridioides difficile* is a graded, microbiota-intrinsic property of healthy human gut communities"

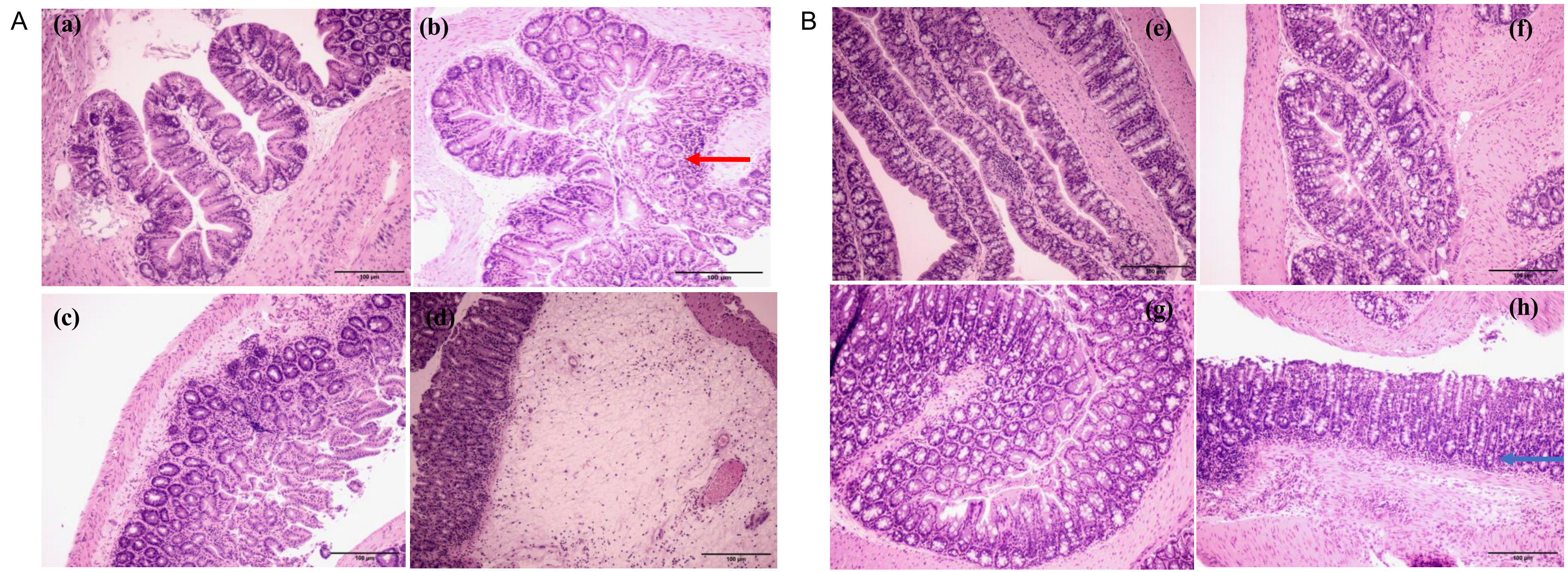


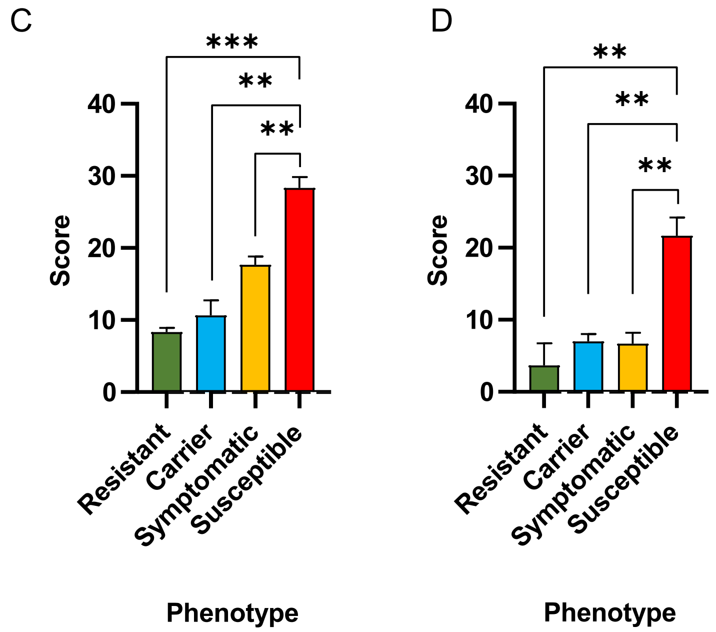


**Supplemental Figure S1. Histopathological analysis of cecum and colon from *C. difficile*-challenged mice.**
**(A)** Representative hematoxylin and eosin–stained sections of cecum from Resistant (a), Carrier (b), Symptomatic (c), and Susceptible (d) mice. **(B)** Representative sections of colon from Resistant (e), Carrier (f), Symptomatic (g), and Susceptible (h) mice. Red arrows in panel b indicate minimal to no inflammation with minimal inflammatory cell infiltrate within the mucosa , and blue arrows in panel h indicate a moderate to marked inflammation with mucosal erosion and inflammatory cell infiltrate of the mucosa and submucosa. **(C)** Histopathological scores in the cecum across phenotypes. **(D)** Histopathological scores in the colon across phenotypes. Scores increased progressively with phenotype severity in both tissues. Data are mean ± SEM. $^{**}p < 0.01$, $^{***}p < 0.001$ (Welch's t-tests).

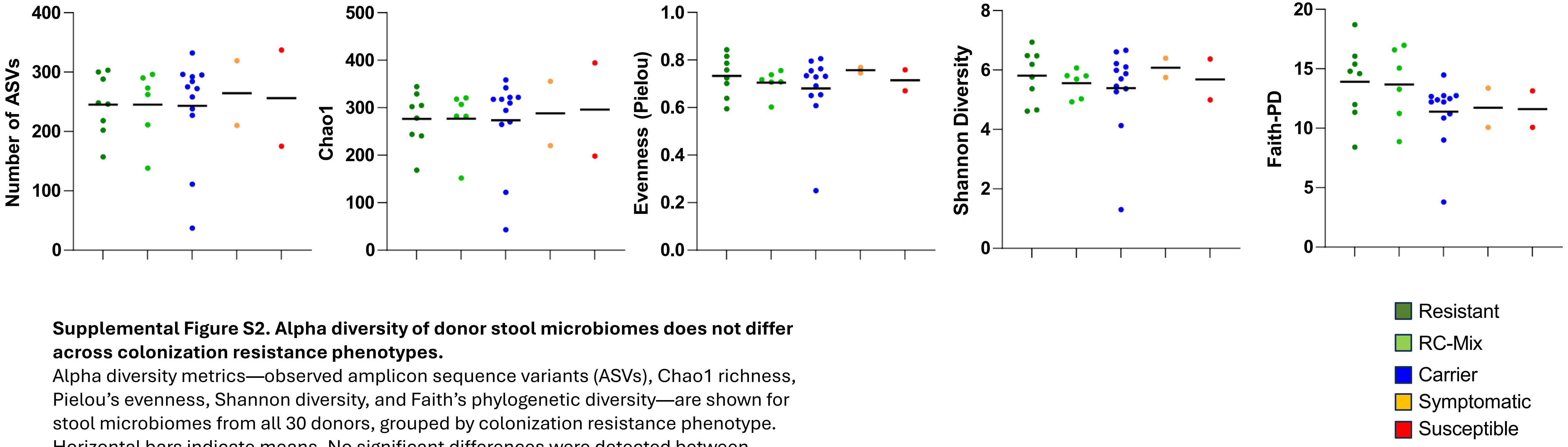


**Supplemental Figure S2. Alpha diversity of donor stool microbiomes does not differ across colonization resistance phenotypes.**
Alpha diversity metrics—observed amplicon sequence variants (ASVs), Chao1 richness, Pielou's evenness, Shannon diversity, and Faith's phylogenetic diversity—are shown for stool microbiomes from all 30 donors, grouped by colonization resistance phenotype. Horizontal bars indicate means. No significant differences were detected between phenotype groups (Kruskal-Wallis Test). Each dot represents one donor.

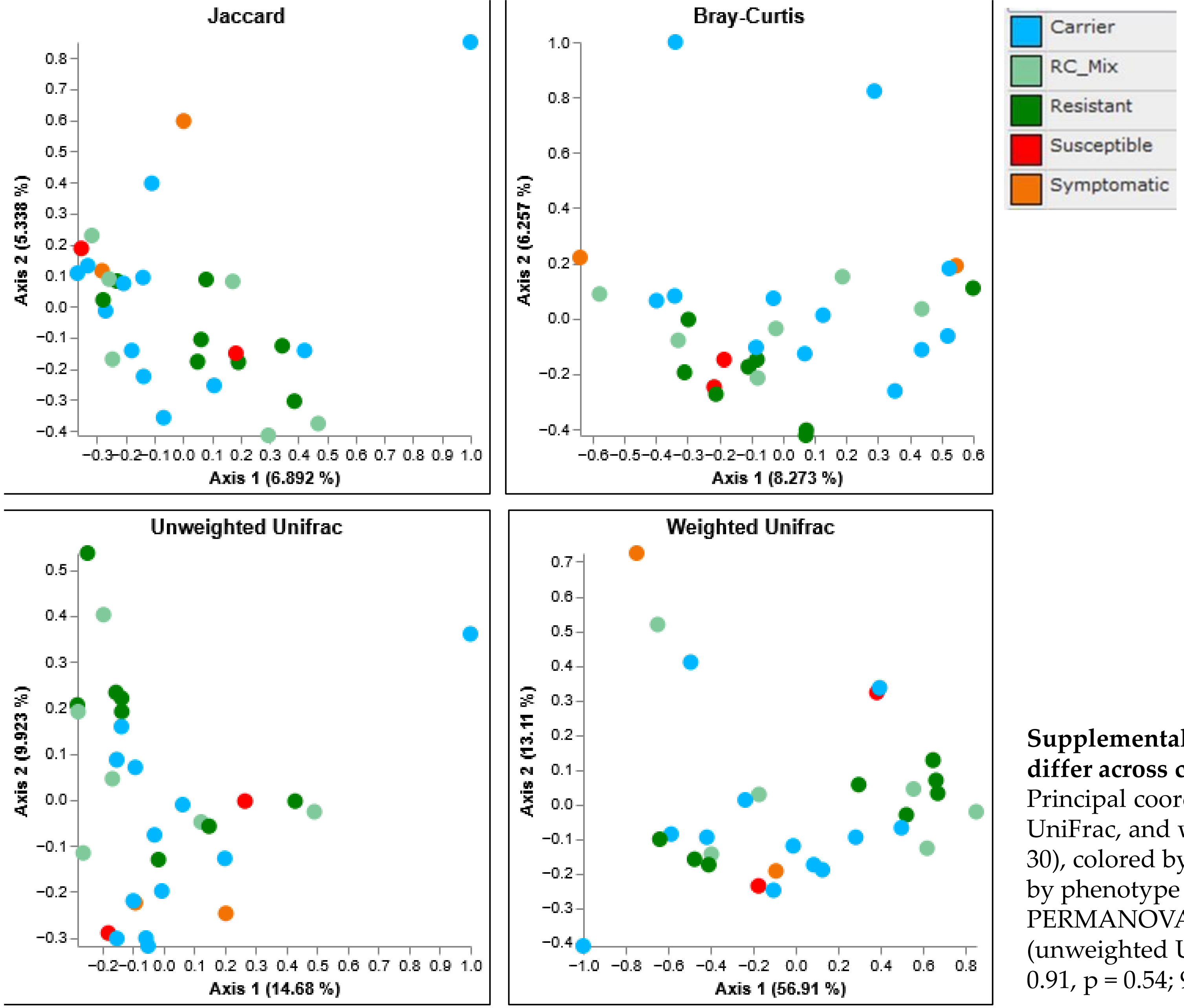


**Supplemental Figure S3. Beta diversity of donor stool microbiomes does not differ across colonization resistance phenotypes.** Principal coordinates analysis (PCoA) of Jaccard, Bray–Curtis, unweighted UniFrac, and weighted UniFrac distances among donor stool microbiomes (N = 30), colored by colonization resistance phenotype. Microbiomes did not cluster by phenotype across any distance metric. Each point represents one donor. PERMANOVA confirmed no significant separation between phenotype groups (unweighted UniFrac: pseudo-F = 0.97, p = 0.516; weighted UniFrac: pseudo-F = 0.91, p = 0.54; 999 permutations; **Supplemental Table S3**).

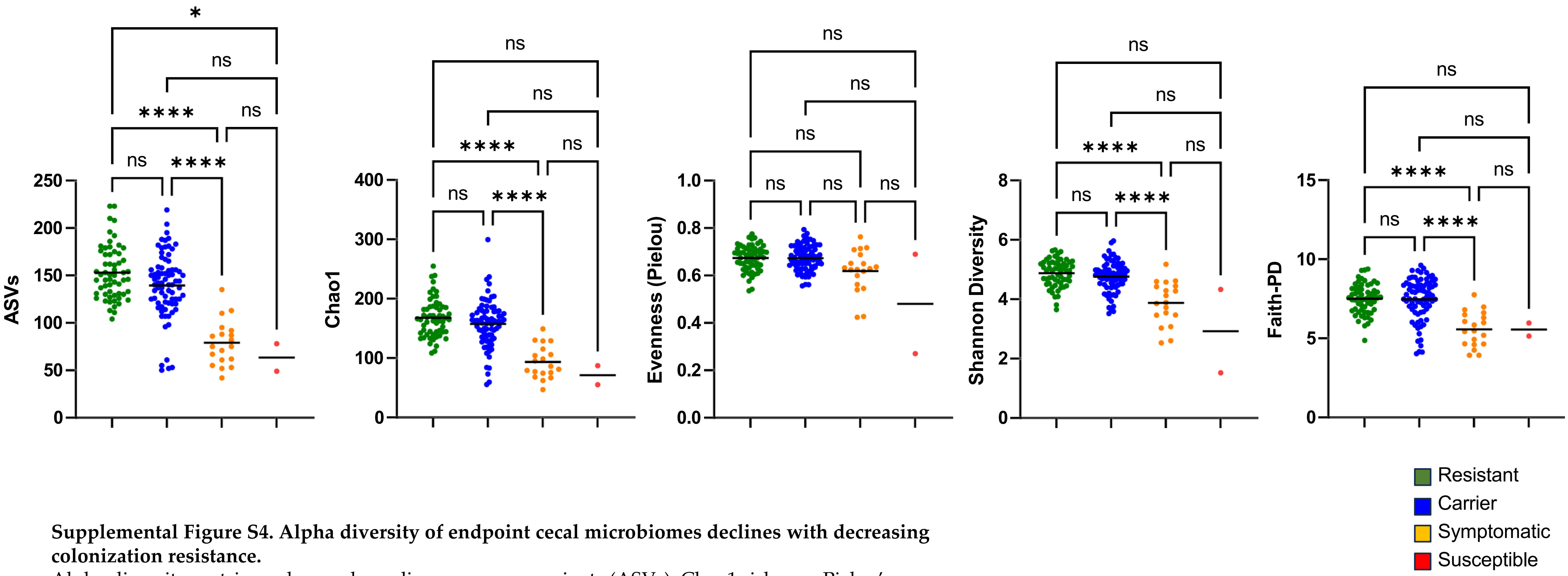


**Supplemental Figure S4. Alpha diversity of endpoint cecal microbiomes declines with decreasing colonization resistance.**
Alpha diversity metrics—observed amplicon sequence variants (ASVs), Chao1 richness, Pielou's evenness, Shannon diversity, and Faith's phylogenetic diversity—are shown for endpoint cecal microbiomes from humanized mice, grouped by colonization resistance phenotype. Cecal contents were collected on day 14 post-challenge for Resistant, Carrier, and Symptomatic mice, and on days 3–4 for Susceptible mice. Horizontal bars indicate means. Richness (observed ASVs and Chao1 richness) and diversity (Shannon diversity and Faith's phylogenetic diversity) were significantly lower in Symptomatic microbiomes than in Resistant and Carrier microbiomes. No significant differences were observed between Resistant and Carrier groups for any metric. Pielou's evenness did not differ significantly across phenotypes. Each dot represents one mouse. *$p < 0.05$, ****$p < 0.0001$; ns, not significant (Kruskal–Wallis test with Dunn's multiple comparisons).

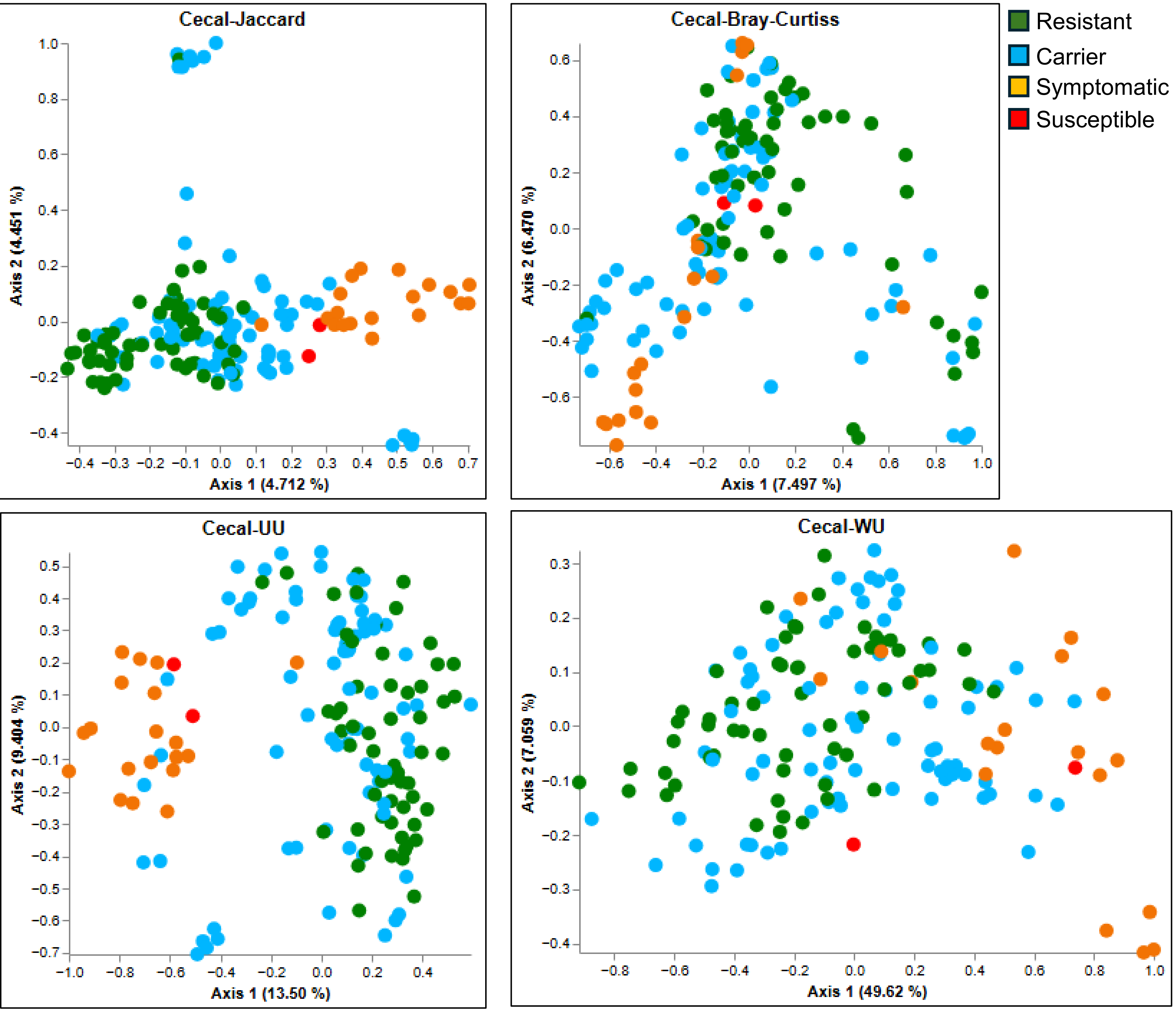


**Supplemental Figure S5. Beta diversity of endpoint cecal microbiomes differs by colonization resistance phenotype.**
Principal coordinates analysis (PCoA) of Jaccard, Bray–Curtis, unweighted UniFrac (UU), and weighted UniFrac (WU) distances among endpoint cecal microbiomes from humanized mice, colored by colonization resistance phenotype. Cecal contents were collected on day 14 post-challenge for Resistant, Carrier, and Symptomatic mice, and on days 3–4 for Susceptible mice. Microbiomes exhibited phenotype-dependent separation across all four distance metrics, with Symptomatic microbiomes showing the greatest divergence from Resistant and Carrier communities. Resistant and Carrier microbiomes showed substantial overlap. Each point represents one mouse. The percentage of variance explained by each axis is indicated on the axis labels. PERMANOVA confirmed significant differences between Resistant, Carrier, and Symptomatic phenotype groups ($p \leq 0.001$; **Supplemental Table S5**).

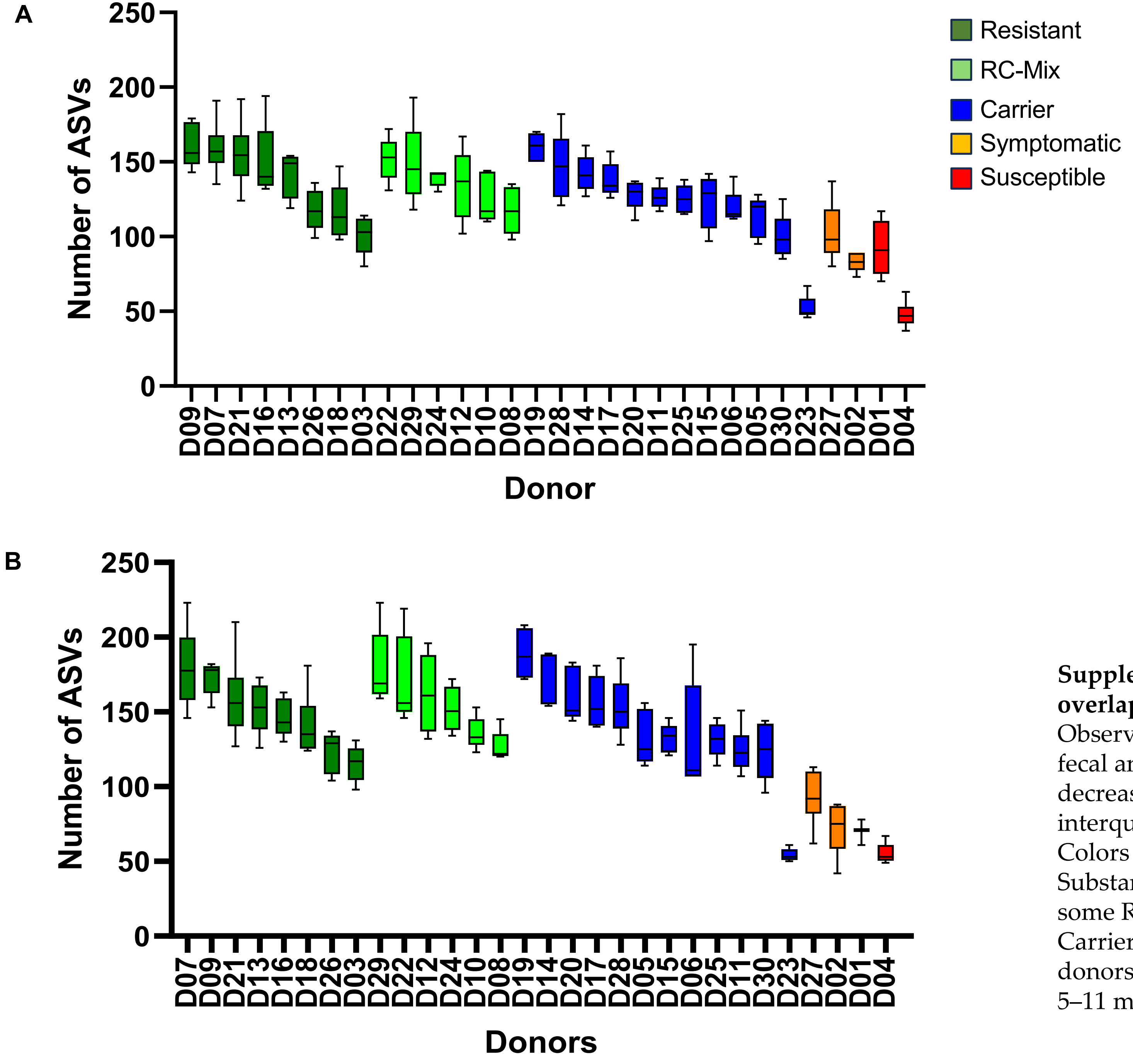


**Supplemental Figure S6. Donor-level variability in microbiome richness overlaps across colonization resistance phenotypes.**
Observed amplicon sequence variants (ASVs) per donor in (A) pre-challenge fecal and (B) endpoint cecal microbiomes of humanized mice, ordered by decreasing mean richness within each phenotype group. Boxes indicate interquartile range, and whiskers indicate minimum and maximum values. Colors indicate the colonization resistance phenotype assigned to each donor. Substantial overlap in richness is observed between phenotype groups, with some Resistant donors exhibiting lower richness than Carrier donors and some Carrier donors exhibiting richness comparable to Symptomatic or Susceptible donors. Each box represents mice colonized from a single donor microbiota (n = 5–11 mice per donor for panel A; n = 3–8 mice per donor for panel B).

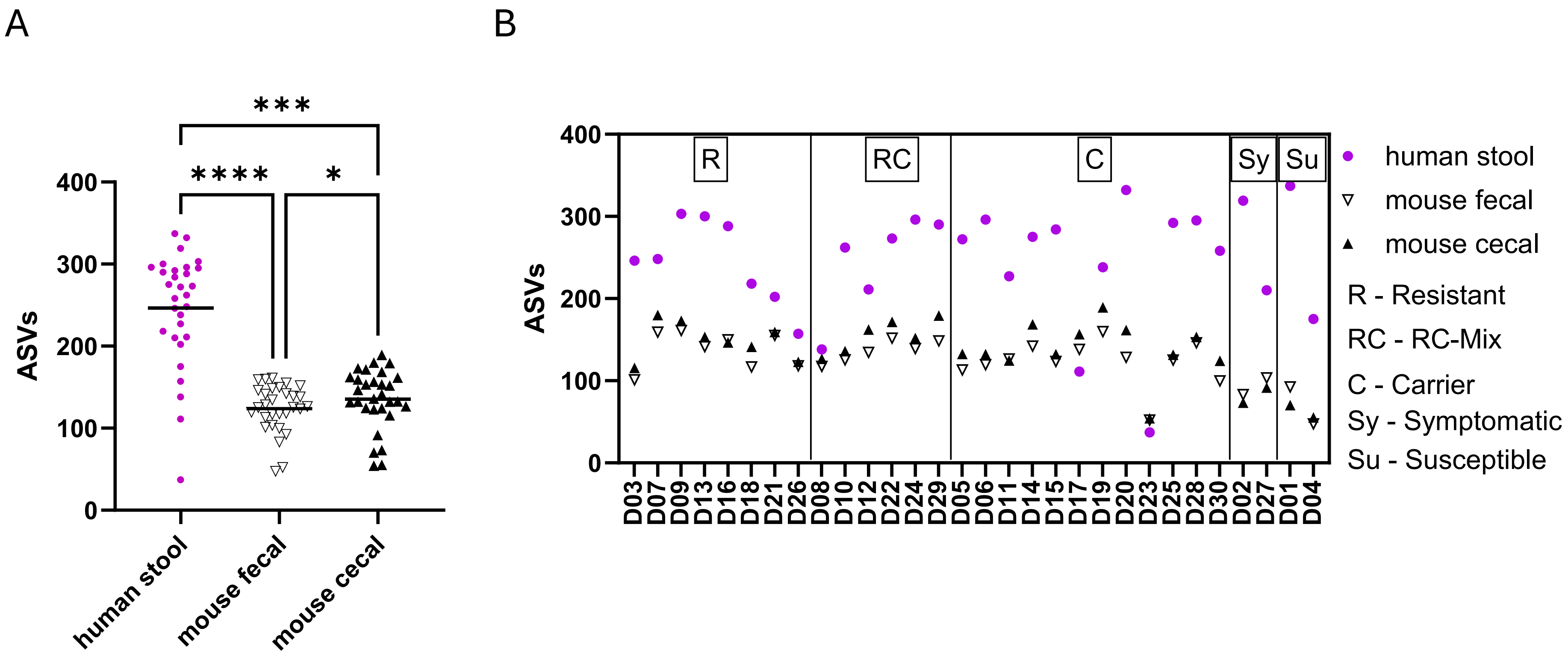


**Supplemental Figure S7. Engraftment of donor microbiota in humanized mice is partial and variable.**
**(A)** Number of amplicon sequence variants (ASVs) detected in human donor stool, mouse pre-challenge fecal microbiomes, and mouse endpoint cecal microbiomes (n = 30 donors per group). ASV counts were significantly lower in mouse microbiomes compared to donor stool. Each point represents one donor (stool) or the mean across recipient mice per donor (fecal and cecal). *$p < 0.05$, ***$p < 0.001$, ****$p < 0.0001$ (Friedman test). **(B)** Donor-wise comparison of ASV counts in human stool (magenta circles), mouse pre-challenge fecal microbiomes (open triangles), and mouse endpoint cecal microbiomes (filled triangles), grouped by colonization resistance phenotype. Vertical lines delineate phenotype groups (R, Resistant; RC, Resistant–Carrier mix; C, Carrier; Sy, Symptomatic; Su, Susceptible). Human stool n = 1 per donor; mouse pre-challenge fecal n = 5–11 mice per donor; mouse endpoint cecal n = 3–8 mice per donor.

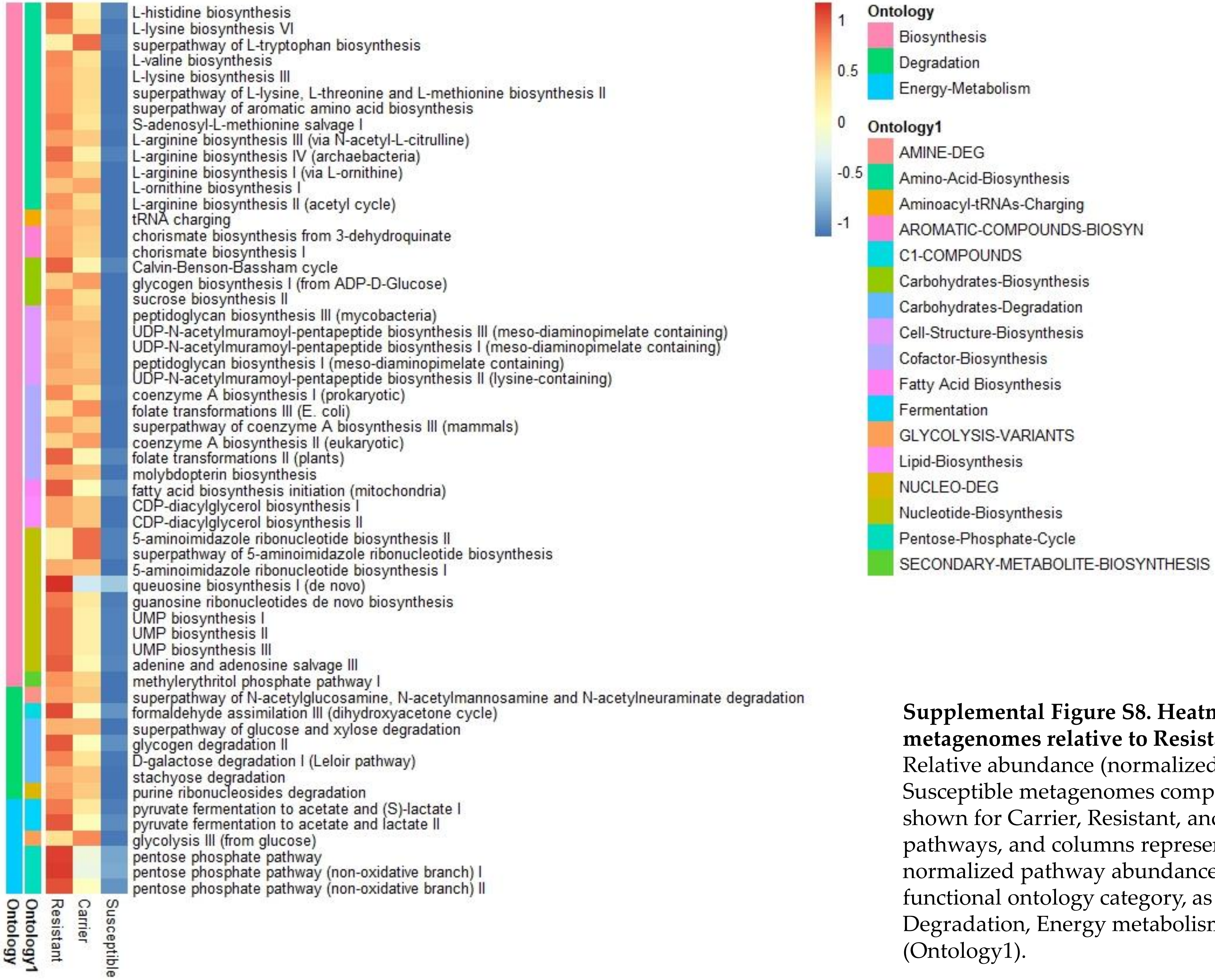


**Supplemental Figure S8. Heatmap of pathways depleted in Susceptible metagenomes relative to Resistant metagenomes.**
Relative abundance (normalized counts) of all 56 pathways significantly depleted in Susceptible metagenomes compared to Resistant metagenomes ($q < 0.05$, MaAsLin3), shown for Carrier, Resistant, and Susceptible groups. Rows represent individual pathways, and columns represent phenotype groups. The color scale indicates normalized pathway abundance (red = high, blue = low). Pathways are grouped by functional ontology category, as indicated in the right panel (Biosynthesis, Degradation, Energy metabolism), and further annotated by pathway subcategory (Ontology1).

**Supplemental Table S1. Mouse health check key**

| Number | Note |
|---|---|
| 0 | Dead |
| 1 | previous severe diarrhea, hunched, eyes starting to close, lethargic/immobile, not willing to move, scruffed fur, shivering *euthanize at this point |
| 2 | severe diarrhea, hobbling, hunching, willing to move, slow, scruffed fur, eyes open, alert |
| 3 | moderate diarrhea, slow but otherwise healthy, alert, no hunching |
| 4 | slight/minor diarrhea, moist stool, clumps, no other symptoms |
| 5 | healthy |

| Gavage from Donor Number | D03 | D07 | D09 | D13 | D16 | D18 | D21 | D26 | D08 | D10 | D12 | D22 | D24 | D29 | D05 | D06 | D11 | D14 | D15 | D17 | D19 | D20 | D23 | D25 | D28 | D30 | D02 | D27 | D01 | D04 |
|---|---|---|---|---|---|---|---|---|---|---|---|---|---|---|---|---|---|---|---|---|---|---|---|---|---|---|---|---|---|---|
| Phenotype of Mouse | C | R | C | R | C | R | C | R | C | C | C | C | C | C | C | C | C | C | C | C | C | C | C | C | C | C | Sy | Sy | Sy | Sy |
| | R | R | R | R | R | R | R | R | C | C | C | C | C | C | C | C | C | C | C | C | C | C | C | C | C | C | Sy | Sy | Sy | Sy |
| | R | R | R | R | R | R | R | R | C | R | C | R | R | C | C | C | C | C | C | C | C | C | C | C | C | C | Sy | Sy | Su | Su |
| | R | R | R | R | R | R | R | R | R | R | R | R | R | R | C | C | C | C | C | C | C | C | C | C | C | C | Sy | Sy | Su | Su |
| | R | R | R | R | R | R | R | R | R | R | R | R | R | R | C | C | C | C | Sy | C | R | C | C | C | R | C | Sy | Sy | Su | Su |
| | | R | | | | R | R | | | | | | | | | | C | | | | | | | | | | | Sy | | Sy |
| | | | | | | | R | | | | | | | | | | C | | | | | | | | | | | Sy | | Sy |
| | | | | | | | R | | | | | | | | | | R | | | | | | | | | | | Sy | | Su |
| | | | | | | | | | | | | | | | | | | | | | | | | | | | | | | Su |
| | | | | | | | | | | | | | | | | | | | | | | | | | | | | | | Su |
| | | | | | | | | | | | | | | | | | | | | | | | | | | | | | | Su |
| Phenotype of Donor Microbiota | R | R | R | R | R | R | R | R | M | M | M | M | M | M | C | C | C | C | C | C | C | C | C | C | C | C | Sy | Sy | Su | Su |

**Supplemental Table S2.** Phenotypes of mice colonized with different human stool microbiota and challenged with *C. difficile* and assignment of phenotypes to donor microbiota based on mice phenotype**.** Donor microbiota was assigned the phenotype exhibited by 80% or more mice. R – Resistant, C –Carrier, Sy – Symptomatic, Su – Susceptible, M – (applicable only to donors) mix of Resistant and Carrier.

**Supplemental Table S3.** PERMANOVA analysis of unweighted and weighted UniFrac distances among donor stool microbiomes.

| | Unweighted | Weighted |
|---|---|---|
| test statistic name | pseudo-F | pseudo-F |
| sample size | 30 | 30 |
| number of groups | 5 | 5 |
| test statistic | 0.973741 | 0.910182 |
| p-value | 0.516 | 0.54 |
| number of permutations | 999 | 999 |

A

| | UU | WU |
|---|---|---|
| sample size | 167 | 167 |
| number of groups | 4 | 4 |
| test statistic | 7.637707 | 12.2308 |
| p-value | 0.001 | 0.001 |
| number of permutations | 999 | 999 |

B

| | | | | UU | | | WU | | |
|---|---|---|---|---|---|---|---|---|---|
| | | Sample size | Permutations | pseudo-F | p-value | **q-value** | pseudo-F | p-value | **q-value** |
| Group 1 | Group 2 | | | | | | | | |
| Carrier | Resistant | 137 | 999 | 3.208994 | 0.001 | 0.0012 | 3.975331 | 0.013 | 0.0156 |
| | Susceptible | 88 | 999 | 7.836546 | 0.001 | 0.0012 | 11.12374 | 0.001 | 0.0015 |
| | Symptomatic | 98 | 999 | 9.165293 | 0.001 | 0.0012 | 14.19335 | 0.001 | 0.0015 |
| Resistant | Susceptible | 69 | 999 | 10.24234 | 0.001 | 0.0012 | 17.36593 | 0.001 | 0.0015 |
| | Symptomatic | 79 | 999 | 12.3879 | 0.001 | 0.0012 | 25.66201 | 0.001 | 0.0015 |
| Susceptible | Symptomatic | 30 | 999 | 3.907825 | 0.002 | 0.002 | 2.638406 | 0.037 | 0.037 |

**Supplemental Table S4.** Pairwise PERMANOVA analysis of unweighted and weighted UniFrac distances among pre-challenge fecal microbiomes.

A

| | UU | WU |
|---|---|---|
| test statistic name | pseudo-F | pseudo-F |
| sample size | 157 | 157 |
| number of groups | 4 | 4 |
| test statistic | 7.349801 | 15.57854 |
| p-value | 0.001 | 0.001 |
| number of permutations | 999 | 999 |

B

| | | | | UU | | | WU | | |
|---|---|---|---|---|---|---|---|---|---|
| | | Sample size | Permutations | pseudo-F | p-value | q-value | pseudo-F | p-value | q-value |
| Group 1 | Group 2 | | | | | | | | |
| Carrier | Resistant | 136 | 999 | 6.255286 | 0.001 | 0.0015 | 8.442017 | 0.001 | 0.002 |
| | Susceptible | 79 | 999 | 2.166858 | 0.014 | 0.0168 | 4.491887 | 0.004 | 0.0048 |
| | Symptomatic | 96 | 999 | 10.39058 | 0.001 | 0.0015 | 21.67681 | 0.001 | 0.002 |
| Resistant | Susceptible | 61 | 999 | 2.810914 | 0.001 | 0.0015 | 7.447416 | 0.002 | 0.003 |
| | Symptomatic | 78 | 999 | 16.12398 | 0.001 | 0.0015 | 43.4244 | 0.001 | 0.002 |
| Susceptible | Symptomatic | 21 | 999 | 1.329471 | 0.23 | 0.23 | 2.18879 | 0.077 | 0.077 |

**Supplemental Table S5.** Pairwise PERMANOVA analysis of unweighted and weighted UniFrac distances among endpoint cecal microbiomes.
